# Epigenetic Regulation of Gene Expression in Cancer: Techniques, Resources, and Analysis

**DOI:** 10.1101/114025

**Authors:** Luciane T Kagohara, Genevieve Stein-O’Brien, Dylan Kelley, Emily Flam, Heather C Wick, Ludmila V Danilova, Hariharan Easwaran, Alexander V Favorov, Jiang Qian, Daria A Gaykalova, Elana J Fertig

## Abstract

Cancer is a complex disease, driven by aberrant activity in numerous signaling pathways in even individual malignant cells. Epigenetic changes are critical mediators of these functional changes that drive and maintain the malignant phenotype. Changes in DNA methylation, histone acetylation and methylation, non-coding RNAs, post-translational modifications are all epigenetic drivers in cancer, independent of changes in the DNA sequence. These epigenetic alterations, once thought to be crucial only for the malignant phenotype maintenance, are now recognized as critical also for disrupting essential pathways that protect the cells from uncontrolled growth, longer survival and establishment in distant sites from the original tissue. In this review, we focus on DNA methylation and chromatin structure in cancer. While associated with cancer, the precise functional role of these alterations is an area of active research using emerging high-throughput approaches and bioinformatics analysis tools. Therefore, this review describes these high-throughput measurement technologies, public domain databases for high-throughput epigenetic data in tumors and model systems, and bioinformatics algorithms for their analysis. Advances in bioinformatics data integration techniques that combine these epigenetic data with genomics data are essential to infer the function of specific epigenetic alterations in cancer, and are therefore also a focus of this review. Future studies using these emerging technologies will elucidate how alterations in the cancer epigenome cooperate with genetic aberrations to cause tumorigenesis initiation and progression. This deeper understanding is essential to future studies that will precisely infer patients’ prognosis and select patients who will be responsive to emerging epigenetic therapies.

## INTRODUCTION

Cancer is a complex disease. The malignant transformation is a multi-step process associated with the accumulation of numerous molecular alterations. These molecular changes impact cellular function within the tumor, its microenvironment, and culminate in the hallmarks of cancer: sustained proliferative signaling, resistance to apoptosis, senescence, angiogenesis, invasion and metastasis, deregulating cellular energetics, avoiding immune destruction, tumor-promoting inflammation, and genome instability and mutation [1]. Numerous genetic alterations (mutations, loss of heterozygosity, deletions, insertions, aneuploidy, etc.) have been associated with carcinogenesis [2] and can sometimes present a clear oncogenic function being considered as cancer driver mutations in such cases [3]. All these genetic alterations ultimately result in aberrant gene expression. However, the landscape of genetic alterations is insufficient to explain the pervasive gene expression changes and alterations to cellular function in cancer [4]. For example, according to the Knudson two-hit hypothesis deletions on both alleles of tumor suppressor genes block the mechanisms in the cell that prevent aberrant cellular growth [5,6]. In many cancers, one of these alterations (or “hits”) is a hereditary or somatic mutation in a tumor suppressor gene and the second “hit” an acquired mutation or copy number loss in the other allele. Nevertheless, sometimes the second genetic “hit” is not observed, but clear changes in gene expression are found and can be explained by the epigenetic alterations [4].

Epigenetic changes are as pervasive in cancer as genetic alterations, and likely are responsible for the hidden source of variation in cancer [4]. Epigenetic changes are heritable traits that impact the phenotype by interfering with gene expression without making changes in the DNA sequence [7,8]. These epigenetic mechanisms include: DNA methylation, chromatin remodeling and, more recently, non-coding RNAs, binding of regulatory proteins, such as CTCF, BORIS or diverse transcription factors also affects chromatin states [9,10].

Epigenetic events can also act concomitantly with other molecular processes in normal states or disease to have persistent gene expression and functional alterations. For example, epigenetic alterations that impact transcription factor binding can explain genome-wide transcription dysregulation that are not associated with genetic variation in cancer [11]. Recent studies have also found that cancer mutations are associated with the chromatin structure of the tissue of origin [12]. These mutations are often enriched in regions of closed chromatin, which are inaccessible to DNA repair genes during replication [12]. Therefore, alterations to chromatin structure may be critical early drivers of cancer.

Epigenetic alterations with functional impacts on gene expression in individual tumor remain key targets of interest because they can be reverted by the use of specific epigenetic therapies [9, 13–15]. Therefore, this review focuses on describing resources for reversible epigenetic events that result or that are directly associated with changes in chromatin organization: DNA methylation and chromatin organization (Figure 1). We describe techniques, model systems, and data resources with high throughput epigenetic data in cancer. We also discuss the emerging bioinformatics techniques for analysis of these data and their integration with high-throughput transcriptional data in cancer. Together, these experimental and computational tools are essential to find the hidden sources of genetic variation in cancer and for precision medicine of novel epigenetic therapies.

**Figure 1.**
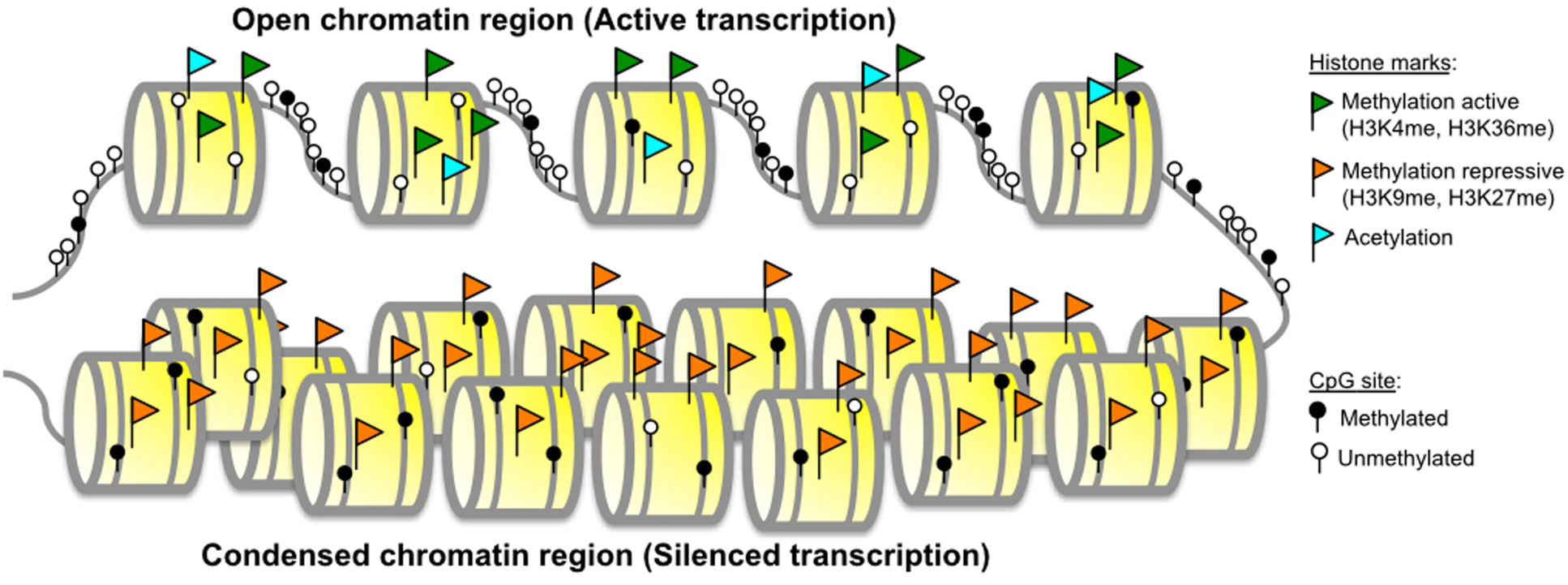
Epigenetic mechanisms of gene expression regulation. Active transcription genes are found in regions where the chromatin conformation is more open, while non-active genes (transcription silencing) are found in regions where the chromatin is more compact. Active chromatin areas are characterized by unmethylated DNA CpG sites, histone acetylation and histone active markers, such as H3K4 and H3K36 methylation. The inverse alterations (methylated CpG sites, histone deacetylation and repressive histone markers) are observed in areas of condensed chromatin, where nucleosomes are positioned closer to each other and DNA accessibility to transcription factors is blocked by the DNA structure and also presence of protein complexes that regulate this compact conformation.

### Reversible epigenetic events

#### DNA Methylation

DNA methylation, the addition of a methyl radical (CH_3_) to the 5-carbon on cytosine residues (5mC) in CpG dinucleotides [16–18], is a normal event with critical roles during different stages of human development, silencing of genome repetitive elements, protection against the integration of viral sequences, genomic imprinting, X chromosome inactivation in females and transcriptional regulation [7,18,19]. In cancer cells, CpG methylation is assessed in tumors relative to that in normal tissues. Removal of CpG methylation specific to cancer cells is referred to as hypomethylation and CpG methylation specific to cancer cells is referred to as aberrant methylation or hypermethylation.

DNA methylation changes to regulatory elements (promoters, insulators, enhancers) with CpG-concentrated regions known as CpG islands (CGI) have been a focus within genome-wide epigenetic studies. These CpGI are extensive sequences (~800 nucleotides in average, range 200-10,000) with higher concentration of CpGs (~10%) and C+G content (>55%) when compared to the rest of the genome (1% of CpG and 42% C+G content) [20,21]. DNA methylation changes outside of CpG island regions or their “CpG shores” are also apparent. The methylation patterns in these shores are highly tissue-specific in normal human samples [22]. The alterations in DNA methylation in cancer extend well beyond CpG island shores [22,23]. These methylation changes have far greater variability than that of normal samples and methylation profiles for some regions distant from CpG islands may be more similar to other normal tissues than the cell of origin [22]. The impact and function of these distant alterations remain poorly understood.

Genome-wide hypomethylation was the first described epigenetic aberration in cancer [24]. Loss of normal DNA methylation levels is associated with genome instability and aneuploidy, reactivation of transposable elements, and loss of imprinting [24,25]. Hypermethylation is associated with gene silencing [26–28], since this event can (1) block transcription directly, by blocking transcription factors binding to their specific sites; or indirectly, by (2) recruitment of protein complexes with high affinity for methylated DNA (methyl-binding domain complexes - MBDs) (Figure 1) [16,17,19,27,29–32]. Therefore, hypermethylation can serve as the second alteration to tumor suppressor genes in Knudson’s two-hit hypothesis [4,5,33–35]. Hypermethylation of gene promoters not only affects the expression of protein coding genes but also constitutes a mechanism to regulate the expression of various noncoding RNAs, some of which have a role in malignant transformation [4].

#### Chromatin modifications

Histone marks are post-translational modifications of core histone proteins that affect chromatin structure [36]. These modifications correlate with open or closed conformations of chromatin, drive differential access of genes to transcription factors and regulatory proteins (Figure 1). Therefore, histone alterations are a facile mechanism for cells to dynamically regulate their gene expression. They are frequently de-regulated in complex diseases, including cancer [4]. Changes in chromatin landscape frequently co-occur with alterations of DNA methylation signals. Histone methylation (20 variants) and acetylation (18 variants) are the most common modifications that mainly target lysine residues of histone tails [37]. Many more modifications include phosphorylation, ubiquitylation, and glycosylation, which spread to other amino acid residues, such as arginine, serine and threonine [38–41]. The critical amino-acids of histone tails most commonly used for diverse modification are found mutated in cancer [42–44]. There are multiple regulatory proteins that write, read and/or erase histone marks [45], many of which also get dysregulated in diseased cells [46]. Because histone marks have stable covalent structures, they can be inherited during cell division and DNA duplication and serve as strong disease markers [47]. Analysis of chromatin structure and its regulatory machinery are key in developing epigenetics therapies.

#### High-throughput Platforms for Epigenetic Analysis

In the current era of high-throughput data, new technologies to measure the genome-wide state of DNA methylation and chromatin structure are actively emerging (Figure 2). Following the history of the field, many measurement platforms were first developed in microarrays and adapted to next generation sequencing technologies. As a result, cancer biologists have access to unprecedented measurement technologies that are able to assess genome-wide DNA methylation and chromatin modifications, accessibility, protein interactions, and binding. Here in this review, we will briefly describe high-throughput approaches for epigenetic mapping from DNA methylation, chromatin modification markers and chromatin structure commonly used in cancer genomics, with details about each measurement technology in Supplemental File 1.

**Figure 2.**
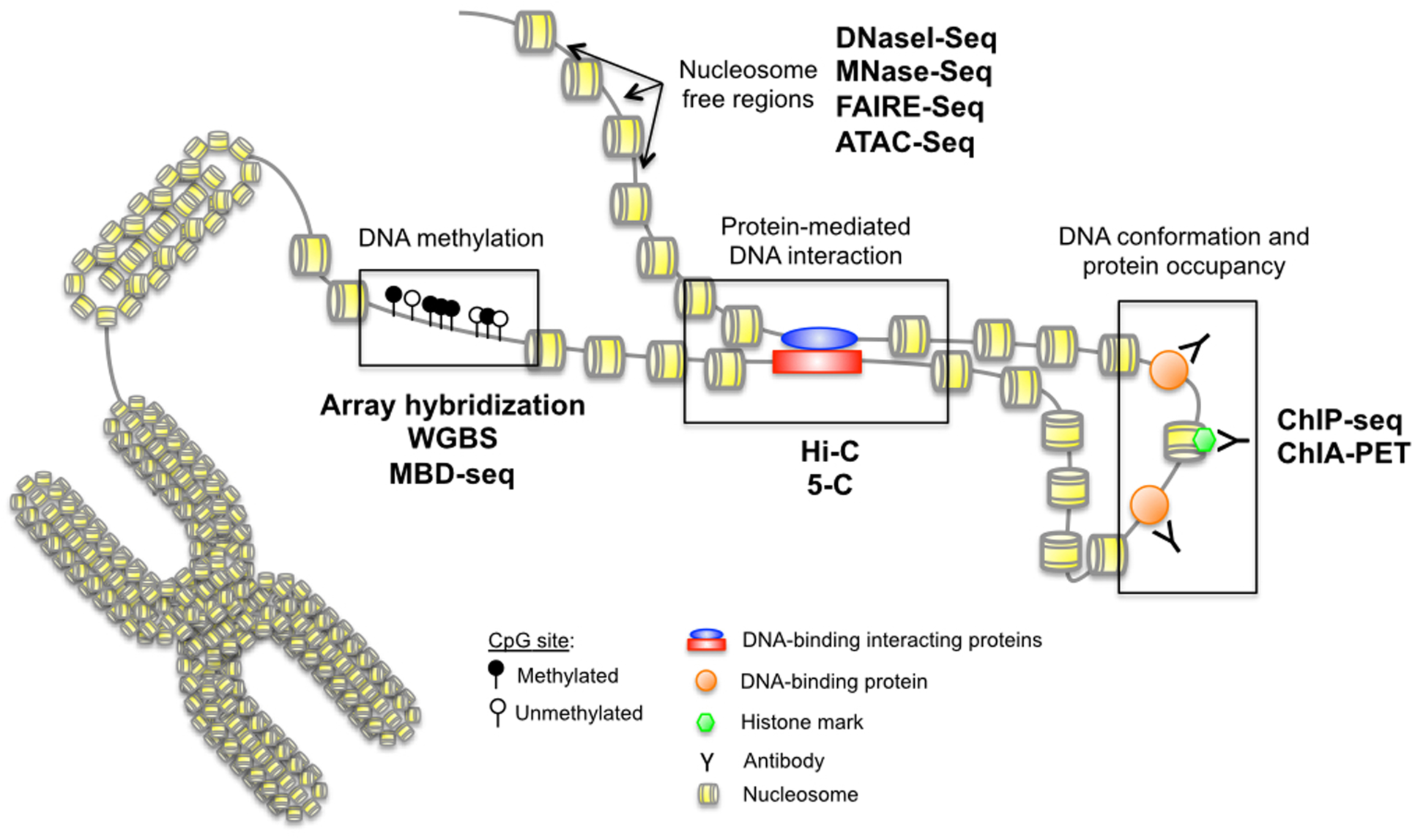
Techniques for epigenetic investigation. There is a wide variety of methods that can be used to characterize epigenetic alterations. Currently, the most common genome-wide approaches allows the identification of nucleosome-free regions (DNaseI-Seq, MNase-Seq, FAIRE-Seq, ATAC-Seq), protein-mediated DNA interaction sites (Hi-C, 5-C), histone marks and DNA binding proteins (ChIP-Seq, ChIA-PET) and DNA methylation (array hybridization, whole genome bisulfite sequencing, MBD-Seq).

### DNA Methylation

Numerous microarray and sequencing-based technologies have been used to measure DNA methylation (Table 1). DNA methylation can be measured on native DNA through recognition of methylated cytosines by antibodies (Methylated DNA immunoprecipitation) or by conjugated Methyl-CpG Binding Proteins (MBP) [51]. The antibodies can also recognize DNA methylation-associated proteins such as MeCP2 with chromatin immunoprecipitation (ChIP)-based technologies, which can be used to estimate DNA methylation. Massively parallel next generation sequencing or arrays to measure DNA methylation-enriched fragments provides quantitative whole-genome evaluation of DNA methylation and allows estimating a high resolution of mCpG sites. False negatives may arise from incomplete binding of the antibodies. Nonetheless, these techniques have strong true positive rates due to the nanomolar binding affinity to symmetrically methylated CpG.

**Table 1.** High-throughput DNA Methylation techniques.

| METHODOLOGY | MeDIP (Methylated DNA immunoprecipitation) | MeCP2-ChIP (Chromatin Immunoprecipitation) | MBP (Methyl-CpG Binding Proteins) | BS (bisulfite sequencing) |
| --- | --- | --- | --- | --- |
| DNA input | Native DNA |  |  | Bisulfite-converted DNA |
| Fragmentation | Sonication |  | Endonuclease |  |
| Enrichment | Antibody (Ab) anti-mCpG | Ab anti-MBP proteins | MBP against mCpG | Bisulfite-converted DNA |
| Control input | Total DNA fraction with no enrichment |  |  | Native DNA |
| Amplification | PCR-based |  |  | PCR-based (no mCpG is amplified as TpG but as CpG) |
| Sequencing | 4-letter based genome |  |  | 3-letter based genome |
| Advantages | High resolution; independence on intermediate steps (e.g.: DNA bisulfite conversion) | Independence on intermediate steps (e.g.: DNA bisulfite conversion) | MBD2 protein has nanomolar affinity for a single symmetrically methylated CpG dinucleotide; MBD2-MBD does not bind unmethylated DNA oligonucleotides to any appreciable extent | Single CpG resolution |
| Disadvantages | Dependence on Ab quality | Lower resolution; dependence on DNA and chromatin integrity | Quantitative methodologies are under development | Dependence on the efficiency of bisulfite conversion step |
| Array-based technologies | MeDIP-chip | ChIP-chip | MBD-chip | Infinium HumanMethylation850 Bead Chip Array from Illumina [Illumina 850K], Human CpG Island Microarray Kit [Agilent], GeneChip Human Promoter 1.0R Arrays |
| Sequence-based technologies | MeDIP-Seq | ChIP-Seq | MBD-Seq | Whole genome bisulfite sequencing (WGBS) |
| References | [48-50] | [51,52] | [53-56] | [57-60] |

Whole-genome DNA methylation signatures can be also performed on DNA converted by bisulfite, which deaminates non-methylated cytosine to uracils to be further recognized as thymidine during sequencing or array-based probe annealing. Notably, during bisulfite conversion methylated cytosines stay unchanged. Both bisulfite converted DNA and untreated input controls are commonly used for arrays (Illumina HumanMethylation Bead Chip Arrays, Agilent Human CpG Island Microarray, and Affymetrix GeneChip Human Promoter 1.0R Arrays). The ratio between methylated and unmethylated signals is proportional to the methylation level of each specific CpG. Bisulfite conversion reduces genome complexity from four nucleotide types to three, except for the few methylated CpG sites, in many cases making alignment to the reference genome or annealing to the particular probe non-unique and introducing errors into the quantification and interpretation. Moreover, the results will depend on the efficiency of bisulfite-conversion step. Therefore, new bioinformatics techniques for preprocessing bisulfite-based data remain a critical challenge preceding the analysis of DNA methylation data in cancer. Comparison of the epigenetic changes across reads enables quantification of intra-tumor variability of epigenetic alterations and of the specificity of epigenetic alterations in a single locus [60]. In spite of the challenges with normalization, these measurements of epigenetic variation within tumors are only possible with bisulfite sequencing techniques. Single cell bisulfite sequencing technologies are emerging to further refine the variability of the epigenetic landscape within tumor samples.

### Chromatin Structure and Interaction

Chromatin is the DNA-protein complex that compacts and protects the genomic DNA within the cellular nucleus and the carrier of epigenetic information, with techniques to measure its structure and interaction domains summarized in Table 2. The structure of chromatin can be evaluated by DNA accessibility to the restriction reagents, such as DNaseI or MNase, or to transposase (ATAC). Chromatin structure also can be evaluated by the binding of DNA to specific proteins (ChIP, FAIRE). The 3D chromatin structure can be defined by close proximity of the distant DNA regions to each other (HiC and ChIA-PET). Just like in case of DNA methylation, fragmented and enriched DNA can be examined with arrays or sequencing to define the whole-genome structure of the chromatin. The binding of DNA by histones or other proteins, such as enzymes or transcription factors decreases the accessibility of such regions to restriction, shredding or transposon insertion.

**Table 2.** High-throughput chromatin organization techniques.

| TYPE OF ANALYSIS | 2D STRUCTURE |  |  |  | 3D STRUCTURE |  |
| --- | --- | --- | --- | --- | --- | --- |
| Methodology | DNase1/MNase (Micrococcal nuclease) | FAIRE (Formaldehyde-Assisted Isolation of Regulatory Elements) | ChIP (Chromatin immunoprecipitation) | ATAC (Assay for Transposase-Accessible Chromatin) | HiC | ChIA-PET (Chromatin Interaction Analysis by Paired-End Tag Sequencing) |
| Sample input | Fresh/frozen tissues |  |  | Fresh cell lines | Fresh tissues |  |
| Fragmentation | Endo- or exonuclease | Sonication | Endo- or exonuclease, sonication | No fragmentation step. Tn5 transposase tagmentation. | Endonuclease |  |
| Enrichment | Size selection | Phenol-chloroform extraction | Antibody (Ab) capture | Specific adapters (tagmentation) | Biotin-streptavidin (fragments are ligated to biotin) | Antibody |
| Control input | Total genomic DNA with no enrichment |  |  |  |  |  |
| Amplification | PCR-based |  |  |  |  |  |
| Sequencing | 4-letter based genome |  |  |  |  |  |
| Advantages | Modest high-resolution | Does not depend on restriction enzyme or buffer composition | Single protein information | Low-input of sample (50,000 cells) | Single protein information |  |
| Disadvantages | High-input of sample (>1 million cells) | Low resolution | Depends on Ab quality | Requires intact nuclei | Depends on Ab quality |  |
| Array-based technologies | DNase1-chip<br>MNase-chip | - | ChIP-chip | - | C3, C4, C5 | - |
| Sequence-based technologies | DNase-Seq<br>MNase-Seq | FAIRE-Seq | ChIP-Seq | ATAC-Seq | C4, C5, HiC | ChIA-PET |
| References | [62-66] | [67] | [68-71] | [72-74] | [75-79] | [80] |

Chromatin analysis reveals whole-genome structure of the chromatin in individual samples and defines regions with open chromatin in functionally active regulatory elements such as promoters, silencers, enhancers, as well as intergenic regions. Due to the modest sequence-dependence of enzymes like DNase1 or MNase, there is a probability of false-positive signals. Public domain databases of histone antibody specificity have been released to provide optimal signal for measurements of histone modifications with ChIP [80].

Other restrictions of chromatin analysis arise from the requirement of high tissue input for most of the procedures and limitation of fresh tissues or cell lines. Both prevent analysis of the majority of primary human tissues. Moreover, individual samples have different cellular properties, and the degree of chromatin digestion by endo-/exonucleases or by sonication must be empirically determined for individual samples to avoid under- and over-digestion. The chromatin integrity and DNA-protein binding strength highly depend on sample preservation. Even upon proper chromatin fragmentation condensed chromatin with repressive marks is under-digested, resulting in presence of longer DNA segments (>900 bp), their poor amplification during library prep for the sequencing, and poor resolution of repressive histone marks mapping. Both ChIP and ChIA-PET require preparation of individual samples for each study protein, and therefore analysis of several proteins can be costly and require high material volume that is not available for most primary cancer samples.

#### Model Systems

Ideally, the extensive epigenetic measurement technologies will be applied to primary tumors samples to measure their epigenetic state. Such extensive profiling of DNA methylation has been performed extensively across measurement technologies. Measuring chromatin, however, proves to be more challenging. Many chromatin assays require large quantities of high quality DNA and the intact chromatin structure. However, tumor samples that are available for profiling are typically small and use preservation techniques that may degrade the quality of the DNA or chromatin structure. Therefore, both *in vitro* and *in vivo* model systems of cancer are essential to determining the epigenetic state of many cancer types (Figure 3). Extending these techniques to humanized PDXs is essential to determine the impact of the immune system to perform preclinical studies correlating the functional role of epigenetics on the efficacy of epigenetic inhibitors and immunotherapy.

**Figure 3.**
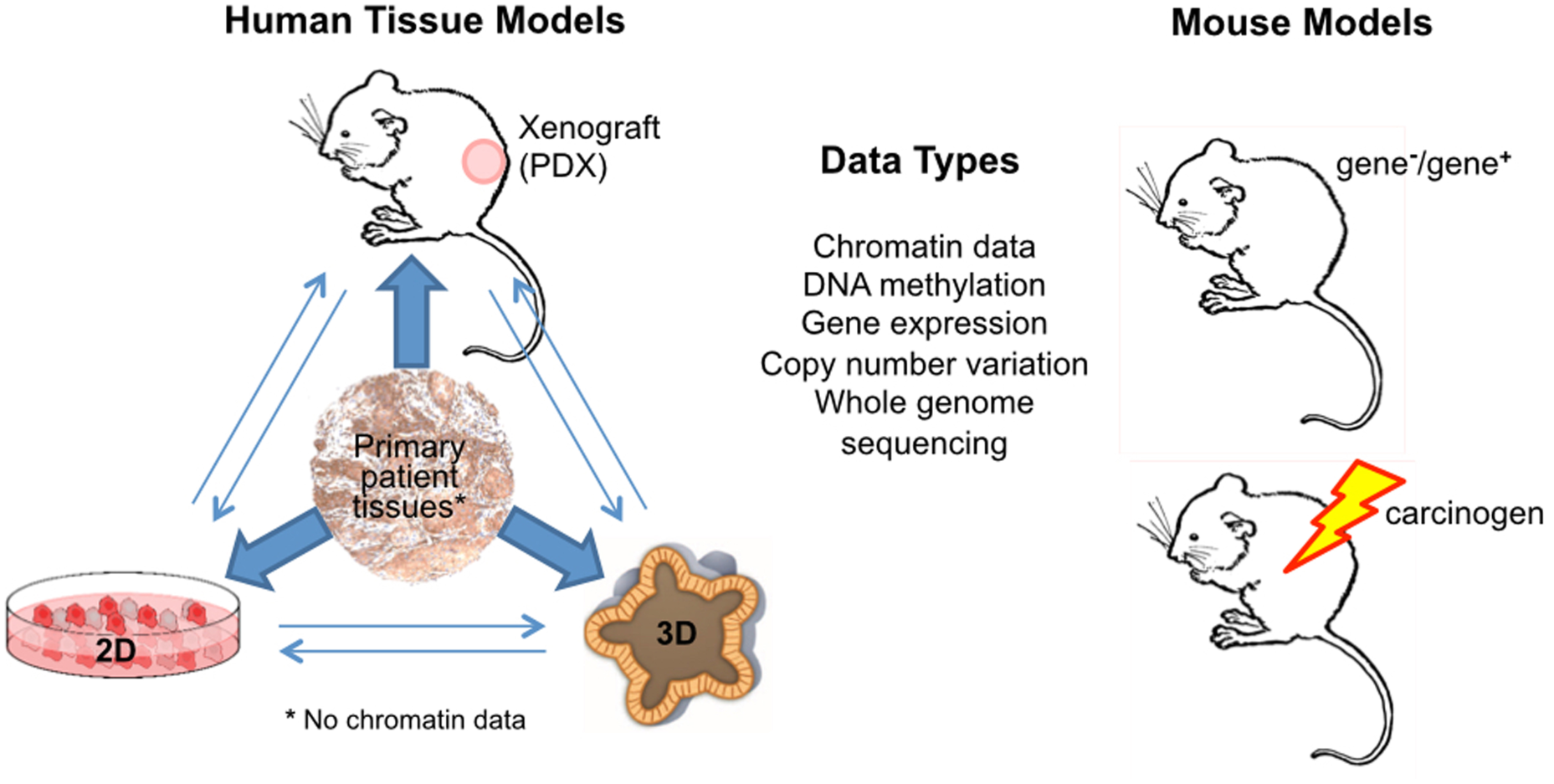
Epigenetic data can be obtained *in vivo* from primary tumor samples, patient derived xenografts (PDX), and mouse models of cancer. While there are no limitations, chromatin structure and interaction data cannot be measured in primary patient samples due to demands of high tissue quantity and quality. All epigenetic data can also be obtained *in vitro* with 2D (cancer cell lines) and 3D (organoids and conditionally reprogrammed cells) culture systems.

#### Epigenetics Data Resources

Numerous international high-throughput genomics databases from thousands of tumor samples and model systems are available in the public domain, reviewed in [81]. However, similar resources for epigenetics in cancer are still more limited. Recent efforts to organize and share large epigenomic datasets have created publicly available resources for discovery and validation in cancer. Cancer-specific resources range in specificity, including large, multi-assay datasets across epigenetics data modalities, such as DNA methylation, histones, and chromatin structure. DNA methylation of 1,001 cancer cell lines was measured with Illumina 450K arrays along with therapeutic sensitivity, copy number, somatic mutations, and gene expression [82] and is freely available from the Gene Expression Omnibus (GEO Series GSE68379). Both the Cancer Genome Atlas (TCGA) and International Cancer Genome Consortium (ICGC) contain microarray measurements of DNA methylation in primary tumors and cancer models. DNA methylation of tumors has been assessed with sequencing technologies in isolated studies from smaller groups, with corresponding data often deposited into sources including GEO [83], dbGAP, and ArrayExpress [84]. However, these sequencing-based DNA methylation data are available for fewer primary tumors than the array-based data from international consortia.

Currently, there are no databases of chromatin structure in primary tumors due to dual challenge of sample quality and quantity described above. Therefore, projects such as ENCODE contain ChIP-seq data of histone marks and chromatin accessibility in numerous cancer cell lines in place of primary tumors [85]. More narrow databases, like FANTOM’s efforts to characterize promoter utilization in different cell types, including 250 cancer cell lines [86]. As with DNA methylation, chromatin measurements in other model organisms have been performed by individual labs. Data of chromatin structure of healthy samples from projects such as the Epigenomics Roadmap [87] have found unanticipated associations with mutations in cancer samples [88]. Therefore, comparison of epigenetics data unrelated to cancer with cancer genomics datasets in other sources may also yield novel insights into the functional impacts of epigenetic regulation in cancer.

In spite of the breadth of public domain chromatin datasets, there is a lack of centralized resources to obtain the wide range of data from numerous studies in a centralized platform. Cistrome is a centralized database of histone modifications [89], which includes Cistrome Cancer to integrate TCGA gene expression data with public ChIP-seq data to determine functional histone modifications in cancer. Another database, EnhancerAtlas (enhanceratlas.org), was specially designed for enhancer analysis and visualization across studies [90]. It contains enhancer annotation for more than 100 cell/tissue types, including both normal and cancer cells. The enhancer annotation was supported by multiple, independent experimental evidences such as chromatin accessibility, histone modifications and eRNAs. Furthermore, the database provides several analytic tools so that the users can compare the enhancer activity across different cell types or connect the enhancers and target genes.

Determining the function of epigenetics alterations in cancer requires integration with genomics data. Ideally, these measurements would be made for the same tumor. Most prominently, the Cancer Genome Atlas (TCGA) and International Cancer Genome Consortium (ICGC) contain microarray measurements of DNA methylation, RNA-sequencing of transcription, and RPPA for protein and phospho-protein states in primary tumors. Availability of high throughput genomics data ENCODE and FANTOM further enables correlation of chromatin features with gene expression to assess the functional changes resulting from epigenetic alterations in tumors or cancer cells and their association with DNA methylation. New resources containing all these sources for data for a wide range of cell lines and primary tissues are essential to establish epigenetic drivers and therapeutic targets in cancer.

#### Bioinformatics Techniques

For all epigenetic data, be it microarray or next generation sequencing, the bioinformatics pipeline follows three major steps: (1) quality control, (2) preprocessing, and (3) analysis [91]. Typically, each of these steps is performed independently for each study and data modality. The bioinformatics techniques for each of these steps are active areas of research, with the maturity of the techniques matching the age of each measurement technology. Once each dataset is understood independently, epigenetic data can be integrated with other cancer genomics data to determine its functional impact. Such robust, integrated techniques are emerging in bioinformatics. Still, further research in data integration is essential to establish epigenetic drivers of carcinogenesis and therapeutic response for precision medicine.

### DNA methylation normalization and analysis

Preprocessing high throughput epigenetic data is critical to obtain accurate results, and techniques for each measurement technology are described in detail in Supplemental File 1. For microarrrays, this entails image processing and normalizing probe intensities. Whole Genome Bisulfite Sequencing (WGBS) techniques require alignment to a reference genome and quantification similar to most second-generation bulk RNA sequencing techniques. As such, most preprocessing pipelines rely on modification of algorithms developed for RNA sequencing. MBD-Seq data adapts peak calling algorithms from ChIP-seq such as MACS [92] to distinguish genomic regions that are methylated. Whereas MBD-seq is non-quantitative, both microarrays and WGBS provide quantitative measurements of the percentage of methylation at each probe or genome coordinate. These estimates can be allele specific for stranded WGBS. Nonetheless, the false positive rate for MBD-Seq is far lower than arrays or WGBS.

As a high-throughput technique, signal from DNA methylation data is often combined with technical artifacts independent of the biological conditions similar to the batch effects established in gene expression studies [93]. These artifacts are most predominant when comparing data across distinct studies, but may also be present within large cohort studies such as TCGA. Visualization tools have been developed to assess these technical artifacts from DNA methylation arrays [94,95] and are an important first step to any analysis of large cohort studies. Many batch correction techniques developed for gene expression microarrays [96,97] have been applied to correct for technical artifacts in DNA methylation arrays [98]. However, care must be taken when adapting these algorithms to maintain the distribution of DNA methylation values. Therefore, normalization techniques that account for this distribution [99,100] and tissue specificity [101] may be better suited to accounting for batch effects in DNA methylation arrays in cancer. Similar to gene expression microarrays, batch correction techniques must be selected in order to preserve signal for the desired analysis [102]. Signaling-based measurements of DNA methylation are not immune to batch effects. However, these have been less studied than DNA methylation arrays. Gene level estimates of DNA methylation from bisulfite sequencing could be corrected with standard expression based techniques, with the same caveats that apply to microarrays. However, these techniques will not extend to locus-specific methylation estimates or DNA methylation from MBD-sequencing. In all cases, obtaining accurate signal requires considering batch effects as part of the experimental design, most especially avoiding perfect confounding between known technical artifacts (e.g., site of tissue source, sampling batch, etc) and experimental conditions [93].

Many analytic tools for DNA methylation data were developed from well-established procedures for gene expression analysis. Accordingly, techniques for detecting robust differences between two or more conditions are the most ubiquitous and reviewed extensively in [103]. This is especially true for work in cancer, in which case-control studies and comparisons of matched tumor and normal tissue from the same individual lend themselves to this type of analysis. Wilcoxon rank-sum tests and *t* tests comparing the methylation status of individual genes between two or more groups are the most basic and commonly employed analysis. To have an impact on expression, DNA methylation changes would ideally be observed over an entire region of the genome. Therefore, bump-hunting algorithms have also been developed to determine differentially that distinguish sample phenotypes [104]. Variably methylated regions inferred with bump-hunting [104] and outlier based analysis algorithms [105] are ideally suited to capture inter-tumor heterogeneity in DNA methylation alterations.

Because MBD-seq data obtains calls of methylated regions, alternative statistical methods either comparing the signal relative to input control or comparing binary calls necessary for its analysis. Techniques to compare peaks across conditions are currently emerging in the literature [106,107], and are primarily divided into linear models comparing peaks similar to DMRs, such as DiffBind [108], hidden Markov Models, such as ChiPDiff [109] and ChromHMM [110], or whole-genome correlations, such as StereoGene [111] and GenometriCorr [112]. For all data platforms, more advanced methods using mixture models, Shannon entropy, logistic regression, non-negative matrix factorization, clustering, feature selection, and correlation to help power between group comparisons.

### Chromatin analysis

The vast majority of techniques to determine chromatin state and binding are derived from ChIP-seq techniques. Therefore, these data use similar preprocessing and differential analysis techniques to those described for MBD-Seq data. Robust standards for quality control and preprocessing were adopted by ENCODE as gold standards for all chromatin based analyses [113]. Determining the chromatin structure specific to cancer cells or cancer subtypes requires differential binding algorithms similar to those described in analysis of MBD-Seq data and reviewed in [106]. In contrast, chromosomal capture methods, i.e. 5C and HiC, produce multiple values representing interaction profiles for each gene. Current bioinformatic tools to analyze this data are summarized in [114] and rapidly developing.

### Determining Functional Impacts of Epigenetic Modifications with Data Integration

Regardless of the data type, biology defines specific relationships between epigenetic regulation and gene or protein expression. Thus, identifying a functional regulatory role from the resulting epigenetic data requires associating the changes with alterations in gene or protein expression. However, technical heterogeneity and confounding from non-biological artifacts, such as batch effects [93], library preparation [115] and antibody quality [116], are problematic within a single data type and can easily grow to be prohibitively complex when integrating across both biological and technical mechanisms [117].

To prevent complications from undesired technical variation, most integrated techniques focus on associating separate analyses of each data type by the co-localization of significant results at the same genetic location, e.g. hypomethylation of the promoter for a given gene is found to be associated with an increase in the gene’s expression level. These methods require matched samples for each set of comparisons a major limitation when dealing with a finite amount of tissue. Additionally, as a given gene in a subtype of cancer is likely to be affected in only a small fraction of individuals, loci based approaches can be unsuccessful in detecting meaningful biological relationships. Thus, techniques, such as OGSA [118] and RTOPPER [119] seek to increase power and biological inference by integrating these univariate differential results over pathway and genes sets. Algorithms integrating these statistics can be adapted to analyze epigenetic regulation of gene expression from distinct genomics datasets from cohorts with similar study design, and need not necessarily have measurements from the same samples in all data modalities. Outlier based approaches for this integration, such as OGSA [118], are best suited to capture inter-tumor heterogeneity of epigenetic pathway regulation.

In contrast to gene-based integration analyses, fully integrated analysis has additional power to identify genes or pathways that are often disrupted by multiple mechanisms but at low frequencies by any one mechanism. Unsupervised algorithms, such as iCluster [120], Amaretto [121] and matrix factorization algorithms [122,123], search for patterns common in these diverse molecular components, regardless of regulatory relationships (Figure 4). Because of their reliance on the concurrence of positive signal, clustering techniques are unable to encode epigenetic silencing a major form of epigenetic regulation on gene expression. The CoGAPS algorithm finds patterns associated with coordinated DNA methylation and expression changes by encoding a distribution that DNA methylation silences gene expression [123]. Similar models of DNA methylation regulation of gene expression are employed to determine genes with a functional impact on cancer subtypes in the MethylMix algorithm [127]. Supervised algorithms overcome this limitation by comparing gene level associations with phenotype in all measurement platforms [124,125] or in several measurement platforms for multiple pathway members [118,119,126]. However, role of DNA methylation in the gene body is not associated with gene expression silencing, and thus more complex to integrate with these techniques. Therefore, further work is needed to develop robust bioinformatics integration algorithms that encode regulatory relationships between genetic and epigenetic alterations as research refines their interrelationship biologically.

**Figure 4.**
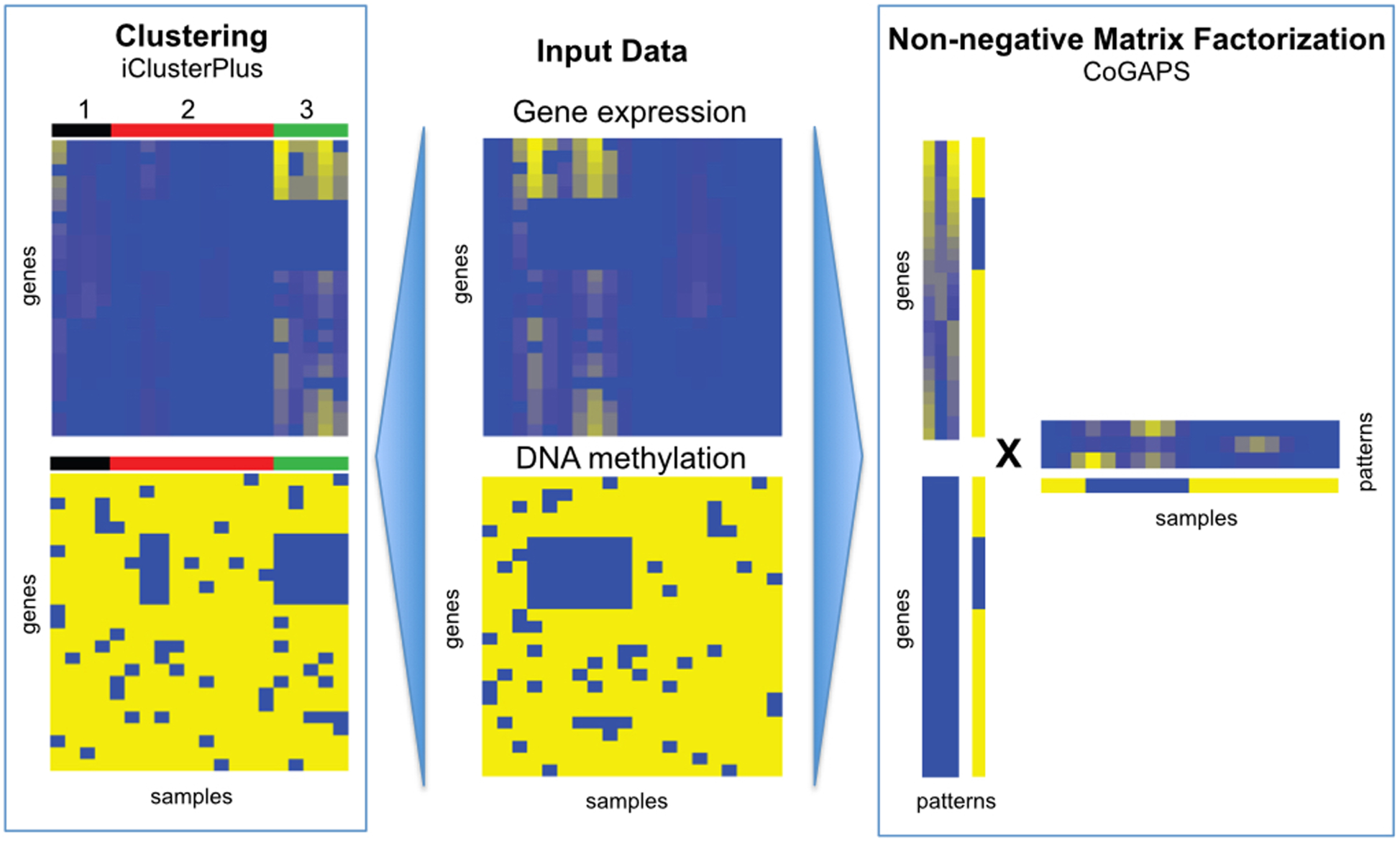
Complete data integration to determine epigenetic regulation of gene expression can be performed for datasets containing both gene expression data (top center) and epigenetic data (bottom center) on the same samples. Clustering-based techniques such as iClusterPlus (left) seek sets of samples that have epigenetic alterations with coordinated gene expression changes. Matrix factorization based techniques such as CoGAPS (right) infer quantitative relationships between epigenetic alterations and gene expression. These algorithms simultaneously quantify the extent of the coordinated alterations in gene expression and epigenetic alterations alterations in each sample. Post-hoc analyses of the clusters in iClusterPlus or extent of epigenetic regulation of gene expression from CoGAPS can determine their functional impact in cancer.

Expanding to the use of biologically driven priors to chromatin data, where different marks or spatial relationships have different regulatory effects, presents additional challenges, and may require adapting techniques developed for dealing with multiple targets in miRNA expression to the chromatin landscape [117,118,125]. Time course methods such as miRDREM have also been developed to determine the timing of activity of miRNA by integrating their expression with that of mRNA targets [128]. These techniques could readily be adapted to determine the functional impacts of chromatin regulation from time course data of cancer development, metastasis, and therapeutic resistance emerging in the literature. Bioinformatics methods are currently being developed that focus on aggregating over epigenetic modifications with similar effects on gene expression, i.e. all repressive or all activating. The ELMER algorithm was developed to incorporate genome-wide maps of enhancers and transcription factors with methylation and expression data to determine epigenetic regulation of transcription factors in cancer [129]. Encoding ways to account to multiple often-conflicting modes of regulation through dysregulation techniques [130] remains a promising avenue for future research.

All the algorithms for integration described above are based upon comparisons of gene- or geneset-level summaries of both the epigenetics and genomics data to associate them with phenotypes in cancer. As is the case for CpG Islands [22], epigenetic alterations in non-coding regions of the genome have critical functional alterations in cancer. In these cases, genomewide associations of epigenetic alterations with the genome, transcriptome, and proteome are essential to determine their functional impact. GenometriCorr [112], which computes the correlation between sets of genomic intervals, can be used to integrate different types of data – such as the location of gene promoters and transcription factor binding sites or other annotation. For a pair of interval-represented datasets, GenometriCorr estimates a variety of correlations that are based on interval overlaps, on relative relative genomic distances, and on absolute genomic distances. GenometriCorr is limited to correlation between binary calls along genome tracks and thus requires an interval calling procedure, e.g. MACS [131] to prepare the interval data for coverage-like (numeric) datasets. In contrast, StereoGene [111] does not require such an identification of intervals. StereoGene uses kernel methods to correlate genome-wide patterns in intensity between two data sets. In this case, epigenetic and genomic profiles may be used as direct inputs for patterns in StereoGene to correlate levels of epigenetic regulation. Both algorithms compute pairwise comparisons between genomic profiles or linear combination of profiles. Therefore, the integration of numerous data types from many samples cannot be computed directly. Instead, these pairwise correlations (or distances) could be inputs to other supervised or unsupervised analysis techniques to infer common epigenetic regulation across cancer samples. Future research into genome-wide coordinate-based integration techniques across multiple samples are essential to determine the full impact of epigenetic alterations on functional genomic alterations in cancer.

## DISCUSSION

Elucidating the relationships between different epigenetic mechanisms and their regulation of gene expression is essential to finding hidden sources of variation in cancer and therapeutic selection. New high-throughput measurement technologies enable unprecedented, quantitative measurements of the epigenetic state in cancers. For DNA methylation, these techniques can be applied readily to both primary tumors and model organisms. Therefore, the functional impact of methylation alterations can be assessed bioinformatically in targeted experiments on model organisms and across sample population. On the other hand, chromatin assays require higher quality and quantity samples that are typical not feasible for preserved tumor samples or biopsies. As a result, chromatin measurements are typically limited to model organisms. In the case of DNA methylation, the epigenetic landscape of cell line models has been shown to vary significantly from that of primary tumors relative to patient-derived xenografts [132]. We anticipate similar discrepancies between model organisms and primary tumors in the chromatin landscape. Thus, advances that adapt chromatin measurement techniques to primary tumor samples are essential to cancer epigenetics.

Databases with epigenetic and genomic data, such as TCGA, ENCODE, and FANTOM are an important step toward achieving this goal. Individually, these large public domain data sets have fueled algorithm development and understanding of epigenetic and tumor based gene expression changes, respectively. While TCGA contains DNA methylation, gene expression, and proteomic data in thousands of primary tumors across cancer types, it lacks chromatin data. Chromatin and transcriptional data are available for numerous cancer cell lines in ENCODE and FANTOM, but these databases lack DNA methylation data and data from primary tumors. However, interactions between DNA methylation and chromatin structure are essential in functional epigenetic regulation. Therefore, it is essential to develop a comprehensive database of matched epigenetic, genetic, and phenotypic data. A comprehensive catalog is especially important for primary tissue samples where cellular heterogeneity compounds the effect of interindividual heterogeneity further obscuring the underlying drivers of disease.

Numerous bioinformatics techniques have been developed to preprocess and analyze single-platform data for DNA methylation and chromatin structure. However, establishing a functional link in cancer requires further identification of epigenetic alterations that are associated with gene expression, protein, and phenotypic changes. Integrating data across measurement platforms is essential to establish these functional relationships. To date, most of these techniques are limited to correlations between genes or common clusters shared across datasets. New integrated bioinformatics techniques are essential to model and distinguish different forms of epigenetic regulation in driving tumor heterogeneity and ultimately cancer. While integrated analyses are emerging, few tools are designed to encode and test these regulatory relationships directly. Determining the true epigenetic regulatory mechanisms and drivers of cancer pathology will be essential for precision medicine with emerging epigenetic therapies.

## KEY POINTS

- Epigenetic alterations compliment genomic alterations during cancer progression and therapeutic response.
- High-throughput measurement technologies can characterize the epigenetic landscape of tumors and model organisms.
- Epigenetic data in large panels of human tumors and cell lines are available from large research consortium.
- Bioinformatics algorithms that integrate epigenetic data with genomics data are essential to determine the function of epigenetic alterations in cancer.

BIOGRAPHICAL NOTES
<u>Luciane Tsukamoto Kagohara</u> is a Postdoctoral Research Fellow in the Department of Oncology Johns Hopkins University SKCCC. She studies the role of aberrant epigenetic and gene expression markers in cancer.
<u>Genevieve Stein-O'Brien</u> is a PhD student at the McKusick-Nathans Institute of Genetic Medicine, Johns Hopkins Medical School. Her research focuses on human development and construction of analytical tools for genetic and epigenetic analysis.
<u>Dylan Kelley</u> is a Researcher in the Department of Otolaryngology, Johns Hopkins University. He is investigating chromatin modification and alternative splicing in transcriptional regulation of head and neck cancer.
<u>Emily Flam</u> is a Researcher in the Department of Otolaryngology, Johns Hopkins University. She is studying the role of enhancers and methylation in transcriptional regulation of head and neck cancer.
<u>Heather Wick</u> is a PhD Candidate at the Institute of Genetic Medicine at Johns Hopkins University School of Medicine. She is studying transcriptional programs and chromosomal remodeling in prostate cancer.
<u>Ludmila Danilova</u> is a Research Associate in the Department of Oncology Johns Hopkins University SKCCC. Her primary interest is in integrative analysis of different types of genomic data.
<u>Hariharan Easwaran</u> is an Assistant Professor in the Department of Oncology Johns Hopkins University SKCCC. He investigates the mechanisms and roles of epigenetic alterations in cancer etiology.
<u>Alexander Favorov</u> is a Research Associate in Johns Hopkins University SKCCC and a Senior Researcher at VIGG RAS and at GosNIIGenetika. He develops statistical approaches to deciphering of biological data.
<u>Jiang Qian</u> is an Associate Professor in the Wilmer Eye Institute, Johns Hopkins University. He works on computational analysis of genetic and epigenetic regulatory networks in various systems.
<u>Daria Gaykalova</u> is an Assistant Professor in the Department of Otolaryngology, Johns Hopkins University. She investigates the role epigenetics in transcriptional regulation of head and neck squamous cell carcinoma progression.
<u>Elana Fertig</u> is an Assistant Professor in the Department of Oncology Johns Hopkins University SKCCC. She develops bioinformatics pattern detection algorithms for epigenetic and genomic data integration in cancer.

## REFERENCES

1. Hanahan D, Weinberg RA. Hallmarks of cancer: the next generation. Cell 2011; 144:646–674

2. Lengauer C, Kinzler KW, Vogelstein B. Genetic instabilities in human cancers. Nature 1998; 396:643–649

3. Vogelstein B, Papadopoulos N, Velculescu VE, et al. Cancer genome landscapes. Science 2013; 339:1546–1558

4. Baylin SB, Jones PA. Epigenetic Determinants of Cancer. Cold Spring Harb. Perspect. Biol. 2016; 8:a019505

5. Berger AH, Knudson AG, Pandolfi PP. A continuum model for tumour suppression. Nature 2011; 476:163–169

6. Knudson AG. Mutation and cancer: statistical study of retinoblastoma. Proc. Natl. Acad. Sci. U. S. A. 1971; 68:820–823

7. Egger G, Liang G, Aparicio A, et al. Epigenetics in human disease and prospects for epigenetic therapy. Nature 2004; 429:457–463

8. Feinberg AP, Tycko B. The history of cancer epigenetics. Nat. Rev. Cancer 2004; 4:143–153

9. Dawson MA, Kouzarides T. Cancer epigenetics: from mechanism to therapy. Cell 2012; 150:12–27

10. Gaykalova D, Vatapalli R, Glazer CA, et al. Dose-Dependent Activation of Putative Oncogene SBSN by BORIS. PLoS ONE 2012; 7:e40389

11. Esteller M. Epigenetics in cancer. N. Engl. J. Med. 2008; 358:1148–1159

12. Polak P, Karlić R, Koren A, et al. Cell-of-origin chromatin organization shapes the mutational landscape of cancer. Nature 2015; 518:360–364

13. Fenaux P, Mufti GJ, Hellstrom-Lindberg E, et al. Efficacy of azacitidine compared with that of conventional care regimens in the treatment of higher-risk myelodysplastic syndromes: a randomised, open-label, phase III study. Lancet Oncol. 2009; 10:223–232

14. Tsai H-C, Li H, Van Neste L, et al. Transient low doses of DNA-demethylating agents exert durable antitumor effects on hematological and epithelial tumor cells. Cancer Cell 2012; 21:430–446

15. Gore SD. New ways to use DNA methyltransferase inhibitors for the treatment of myelodysplastic syndrome. Hematol. Am. Soc. Hematol. Educ. Program 2011; 2011:550–555

16. Jones PA, Laird PW. Cancer epigenetics comes of age. Nat. Genet. 1999; 21:163–167

17. Plass C. Cancer epigenomics. Hum. Mol. Genet. 2002; 11:2479–2488

18. Laird PW. The power and the promise of DNA methylation markers. Nat. Rev. Cancer 2003; 3:253–266

19. Robertson KD. DNA methylation and human disease. Nat. Rev. Genet. 2005; 6:597–610

20. Gardiner-Garden M, Frommer M. CpG islands in vertebrate genomes. J. Mol. Biol. 1987; 196:261–282

21. Deaton AM, Bird A. CpG islands and the regulation of transcription. Genes Dev. 2011; 25:1010–1022

22. Irizarry RA, Ladd-Acosta C, Wen B, et al. The human colon cancer methylome shows similar hypo- and hypermethylation at conserved tissue-specific CpG island shores. Nat. Genet. 2009; 41:178–186

23. Hellman A, Chess A. Gene Body-Specific Methylation on the Active X Chromosome. Science 2007; 315:1141–1143

24. Feinberg AP, Vogelstein B. Hypomethylation distinguishes genes of some human cancers from their normal counterparts. Nature 1983; 301:89–92

25. Kagohara LT, Schussel JL, Subbannayya T, et al. Global and gene-specific DNA methylation pattern discriminates cholecystitis from gallbladder cancer patients in Chile. Future Oncol. Lond. Engl. 2015; 11:233–249

26. Bird AP. DNA methylation--how important in gene control? Nature 1984; 307:503–504

27. Sidransky D. Emerging molecular markers of cancer. Nat. Rev. Cancer 2002; 2:210–219

28. Baylin SB, Ohm JE. Epigenetic gene silencing in cancer - a mechanism for early oncogenic pathway addiction? Nat. Rev. Cancer 2006; 6:107–116

29. Robertson KD, Jones PA. DNA methylation: past, present and future directions. Carcinogenesis 2000; 21:461–467

30. Baylin SB, Esteller M, Rountree MR, et al. Aberrant patterns of DNA methylation, chromatin formation and gene expression in cancer. Hum. Mol. Genet. 2001; 10:687–692

31. Jones PA, Baylin SB. The fundamental role of epigenetic events in cancer. Nat. Rev. Genet. 2002; 3:415–428

32. Worm J, Guldberg P. DNA methylation: an epigenetic pathway to cancer and a promising target for anticancer therapy. J. Oral Pathol. Med. Off. Publ. Int. Assoc. Oral Pathol. Am. Acad. Oral Pathol. 2002; 31:443–449

33. Grady WM, Willis J, Guilford PJ, et al. Methylation of the CDH1 promoter as the second genetic hit in hereditary diffuse gastric cancer. Nat. Genet. 2000; 26:16–17

34. Esteller M, Fraga MF, Guo M, et al. DNA methylation patterns in hereditary human cancers mimic sporadic tumorigenesis. Hum. Mol. Genet. 2001; 10:3001–3007

35. Merlo A, Herman JG, Mao L, et al. 5’ CpG island methylation is associated with transcriptional silencing of the tumour suppressor p16/CDKN2/MTS1 in human cancers. Nat. Med. 1995; 1:686–692

36. Barth TK, Imhof A. Fast signals and slow marks: the dynamics of histone modifications. Trends Biochem. Sci. 2010; 35:618–626

37. Wang Z, Schones DE, Zhao K. Characterization of human epigenomes. Curr. Opin. Genet. Dev. 2009; 19:127–134

38. Bhaumik SR, Smith E, Shilatifard A. Covalent modifications of histones during development and disease pathogenesis. Nat. Struct. Mol. Biol. 2007; 14:1008–1016

39. Turner BM. Cellular memory and the histone code. Cell 2002; 111:285–291

40. Strahl BD, Allis CD. The language of covalent histone modifications. Nature 2000; 403:41–45

41. Sun Z-W, Allis CD. Ubiquitination of histone H2B regulates H3 methylation and gene silencing in yeast. Nature 2002; 418:104–108

42. Chan K-M, Fang D, Gan H, et al. The histone H3.3K27M mutation in pediatric glioma reprograms H3K27 methylation and gene expression. Genes Dev. 2013; 27:985–990

43. Papillon-Cavanagh S, Lu C, Gayden T, et al. Impaired H3K36 methylation defines a subset of head and neck squamous cell carcinomas. Nat. Genet. 2017; 49:180–185

44. Lu C, Jain SU, Hoelper D, et al. Histone H3K36 mutations promote sarcomagenesis through altered histone methylation landscape. Science 2016; 352:844–849

45. Tarakhovsky A. Tools and landscapes of epigenetics. Nat. Immunol. 2010; 11:565–568

46. Rodríguez-Paredes M, Esteller M. Cancer epigenetics reaches mainstream oncology. Nat. Med. 2011; 330–339

47. Judes G, Dagdemir A, Karsli-Ceppioglu S, et al. H3K4 acetylation, H3K9 acetylation and H3K27 methylation in breast tumor molecular subtypes. Epigenomics 2016; 8:909–924

48. Weber M, Davies JJ, Wittig D, et al. Chromosome-wide and promoter-specific analyses identify sites of differential DNA methylation in normal and transformed human cells. Nat. Genet. 2005; 37:853–862

49. Vucic EA, Wilson IM, Campbell JM, et al. Methylation analysis by DNA immunoprecipitation (MeDIP). Methods Mol. Biol. Clifton NJ 2009; 556:141–153

50. Mohn F, Weber M, Schübeler D, et al. Methylated DNA immunoprecipitation (MeDIP). Methods Mol. Biol. Clifton NJ 2009; 507:55–64

51. Gabel HW, Kinde B, Stroud H, et al. Disruption of DNA-methylation-dependent long gene repression in Rett syndrome. Nature 2015; 522:89–93

52. Ballestar E, Paz MF, Valle L, et al. Methyl-CpG binding proteins identify novel sites of epigenetic inactivation in human cancer. EMBO J. 2003; 22:6335–6345

53. Serre D, Lee BH, Ting AH. MBD-isolated Genome Sequencing provides a high-throughput and comprehensive survey of DNA methylation in the human genome. Nucleic Acids Res. 2010; 38:391–399

54. Yegnasubramanian S, Lin X, Haffner MC, et al. Combination of methylated-DNA precipitation and methylation-sensitive restriction enzymes (COMPARE-MS) for the rapid, sensitive and quantitative detection of DNA methylation. Nucleic Acids Res. 2006; 34:e19

55. Yegnasubramanian S, Wu Z, Haffner MC, et al. Chromosome-wide mapping of DNA methylation patterns in normal and malignant prostate cells reveals pervasive methylation of gene-associated and conserved intergenic sequences. BMC Genomics 2011; 12:313

56. Laird PW. Principles and challenges of genomewide DNA methylation analysis. Nat. Rev. Genet. 2010; 11:191–203

57. Kurdyukov S, Bullock M. DNA Methylation Analysis: Choosing the Right Method. Biology 2016; 5:3

58. Adorján P, Distler J, Lipscher E, et al. Tumour class prediction and discovery by microarray-based DNA methylation analysis. Nucleic Acids Res. 2002; 30:e21

59. Gitan RS, Shi H, Chen C-M, et al. Methylation-specific oligonucleotide microarray: a new potential for high-throughput methylation analysis. Genome Res. 2002; 12:158–164

60. Landan G, Cohen NM, Mukamel Z, et al. Epigenetic polymorphism and the stochastic formation of differentially methylated regions in normal and cancerous tissues. Nat. Genet. 2012; 44:1207–1214

61. Wu C, Wong YC, Elgin SC. The chromatin structure of specific genes: II. Disruption of chromatin structure during gene activity. Cell 1979; 16:807–814

62. Wu HM, Dattagupta N, Hogan M, et al. Unfolding of nucleosomes by ethidium binding. Biochemistry (Mosc.) 1980; 19:626–634

63. Song L, Crawford GE. DNase-seq: a high-resolution technique for mapping active gene regulatory elements across the genome from mammalian cells. Cold Spring Harb. Protoc. 2010; 2010:pdb.prot5384

64. Boyle AP, Davis S, Shulha HP, et al. High-resolution mapping and characterization of open chromatin across the genome. Cell 2008; 132:311–322

65. Meyer CA, Liu XS. Identifying and mitigating bias in next-generation sequencing methods for chromatin biology. Nat. Rev. Genet. 2014; 15:709–721

66. Giresi PG, Kim J, McDaniell RM, et al. FAIRE (Formaldehyde-Assisted Isolation of Regulatory Elements) isolates active regulatory elements from human chromatin. Genome Res. 2007; 17:877–885

67. Buenrostro JD, Giresi PG, Zaba LC, et al. Transposition of native chromatin for fast and sensitive epigenomic profiling of open chromatin, DNA-binding proteins and nucleosome position. Nat. Methods 2013; 10:1213–1218

68. Buenrostro JD, Wu B, Litzenburger UM, et al. Single-cell chromatin accessibility reveals principles of regulatory variation. Nature 2015; 523:486–490

69. Pott S, Lieb JD. Single-cell ATAC-seq: strength in numbers. Genome Biol. 2015; 16:172

70. Ho JWK, Bishop E, Karchenko PV, et al. ChIP-chip versus ChIP-seq: lessons for experimental design and data analysis. BMC Genomics 2011; 12:134

71. O’Geen H, Echipare L, Farnham PJ. Using ChIP-seq technology to generate high-resolution profiles of histone modifications. Methods Mol. Biol. Clifton NJ 2011; 791:265–286

72. Schmidt D, Wilson MD, Spyrou C, et al. ChIP-seq: using high-throughput sequencing to discover protein-DNA interactions. Methods San Diego Calif 2009; 48:240–248

73. Bentley DR, Balasubramanian S, Swerdlow HP, et al. Accurate whole human genome sequencing using reversible terminator chemistry. Nature 2008; 456:53–59

74. Dekker J, Rippe K, Dekker M, et al. Capturing chromosome conformation. Science 2002; 295:1306–1311

75. Barutcu AR, Fritz AJ, Zaidi SK, et al. C-ing the Genome: A Compendium of Chromosome Conformation Capture Methods to Study Higher-Order Chromatin Organization: CHROMOSOME CONFORMATION CAPTURE METHODS. J. Cell. Physiol. 2016; 231:31–35

76. Simonis M, Klous P, Splinter E, et al. Nuclear organization of active and inactive chromatin domains uncovered by chromosome conformation capture-on-chip (4C). Nat. Genet. 2006; 38:1348–1354

77. Dostie J, Richmond TA, Arnaout RA, et al. Chromosome Conformation Capture Carbon Copy (5C): a massively parallel solution for mapping interactions between genomic elements. Genome Res. 2006; 16:1299–1309

78. Lieberman-Aiden E, van Berkum NL, Williams L, et al. Comprehensive mapping of long range interactions reveals folding principles of the human genome. Science 2009; 326:289–293

79. Fullwood MJ, Wei C-L, Liu ET, et al. Next-generation DNA sequencing of paired-end tags (PET) for transcriptome and genome analyses. Genome Res. 2009; 19:521–532

80. Rothbart SB, Dickson BM, Raab JR, et al. An Interactive Database for the Assessment of Histone Antibody Specificity. Mol. Cell 2015; 59:502–511

81. Kannan L, Ramos M, Re A, et al. Public data and open source tools for multi-assay genomic investigation of disease. Brief. Bioinform. 2016; 17:603–615

82. Iorio F, Knijnenburg TA, Vis DJ, et al. A Landscape of Pharmacogenomic Interactions in Cancer. Cell 2016; 166:740–754

83. Edgar R, Domrachev M, Lash AE. Gene Expression Omnibus: NCBI gene expression and hybridization array data repository. Nucleic Acids Res. 2002; 30:207–210

84. N K, E H, M K, et al. ArrayExpress update--simplifying data submissions. Nucleic Acids Res. 2015; 43:D1113–6

85. Consortium TEP, data analysis coordination O coordination, data production D production leads, et al. An integrated encyclopedia of DNA elements in the human genome. Nature 2012; 488:57–74

86. Andersson R, Gebhard C, Miguel-Escalada I, et al. An atlas of active enhancers across human cell types and tissues. Nature 2014; 507:455–461

87. Kundaje A, Meuleman W, Ernst J, et al. Integrative analysis of 111 reference human epigenomes. Nature 2015; 518:317–330

88. Chen Y, Sprung R, Tang Y, et al. Lysine propionylation and butyrylation are novel post-translational modifications in histones. Mol. Cell. Proteomics MCP 2007; 6:812–819

89. Liu T, Ortiz JA, Taing L, et al. Cistrome: an integrative platform for transcriptional regulation studies. Genome Biol. 2011; 12:R83

90. Gao T, He B, Liu S, et al. EnhancerAtlas: a resource for enhancer annotation and analysis in 105 human cell/tissue types. Bioinformatics 2016; 32:3543–3551

91. Fertig EJ, Slebos R, Chung CH. Application of genomic and proteomic technologies in biomarker discovery. Am. Soc. Clin. Oncol. Educ. Book Am. Soc. Clin. Oncol. Meet. 2012; 377–382

92. Feng J, Liu T, Qin B, et al. Identifying ChIP-seq enrichment using MACS. Nat. Protoc. 2012; 7:1728–1740

93. Leek JT, Scharpf RB, Bravo HC, et al. Tackling the widespread and critical impact of batch effects in high-throughput data. Nat. Rev. Genet. 2010; 11:733–739

94. mBatch.

95. Fortin J-P, Fertig E, Hansen K. shinyMethyl: interactive quality control of Illumina 450k DNA methylation arrays in R. F1000Research 2014;

96. Johnson WE, Li C, Rabinovic A. Adjusting batch effects in microarray expression data using empirical Bayes methods. Biostatistics 2007; 8:118–127

97. Leek JT, Storey JD. Capturing Heterogeneity in Gene Expression Studies by Surrogate Variable Analysis. PLoS Genet. 2007; 3:e161

98. Maksimovic J, Gagnon-Bartsch JA, Speed TP, et al. Removing unwanted variation in a differential methylation analysis of Illumina HumanMethylation450 array data. Nucleic Acids Res. 2015; 43:e106–e106

99. Fortin J-P, Triche TJ, Hansen KD. Preprocessing, normalization and integration of the Illumina HumanMethylationEPIC array with minfi. Bioinformatics 2016; btw691

100. Fortin J-P, Labbe A, Lemire M, et al. Functional normalization of 450k methylation array data improves replication in large cancer studies. Genome Biol. 2014; 15:

101. Oros Klein K, Grinek S, Bernatsky S, et al. funtooNorm: an R package for normalization of DNA methylation data when there are multiple cell or tissue types. Bioinformatics 2016; 32:593–595

102. Parker HS, Leek JT, Favorov AV, et al. Preserving biological heterogeneity with a permuted surrogate variable analysis for genomics batch correction. Bioinformatics 2014; 30:2757–2763

103. Bock C. Analysing and interpreting DNA methylation data. Nat. Rev. Genet. 2012; 13:705–719

104. Jaffe AE, Feinberg AP, Irizarry RA, et al. Significance analysis and statistical dissection of variably methylated regions. Biostat. Oxf. Engl. 2012; 13:166–178

105. Gaykalova DA, Vatapalli R, Wei Y, et al. Outlier Analysis Defines Zinc Finger Gene Family DNA Methylation in Tumors and Saliva of Head and Neck Cancer Patients. PLOS ONE 2015; 10:e0142148

106. Steinhauser S, Kurzawa N, Eils R, et al. A comprehensive comparison of tools for differential ChIP-seq analysis. Brief. Bioinform. 2016; bbv110

107. Pepke S, Wold B, Mortazavi A. Computation for ChIP-seq and RNA-seq studies. Nat. Methods 2009; 6:S22–S32

108. Ross-Innes CS, Stark R, Teschendorff AE, et al. Differential oestrogen receptor binding is associated with clinical outcome in breast cancer. Nature 2012;

109. Xu H, Sung W. Identifying Differential Histone Modification Sites from ChIP-seq Data. Gener. Microarray Bioinforma. 2012; 802:293–303

110. Ernst J, Kellis M. ChromHMM: automating chromatin-state discovery and characterization. Nat. Methods 2012; 9:215–216

111. Stravrovskaya ED, Favorov AV, Niranjan T, et al. StereoGene: Rapid Estimation of Genomewide Correlation of Continuous or Interval Feature Data. Bioarxiv

112. Favorov A, Mularoni L, Cope LM, et al. Exploring Massive, Genome Scale Datasets with the GenometriCorr Package. PLoS Comput. Biol. 2012; 8:e1002529

113. Landt SG, Marinov GK, Kundaje A, et al. ChIP-seq guidelines and practices of the ENCODE and modENCODE consortia. Genome Res. 2012; 22:1813–1831

114. Schmitt AD, Hu M, Ren B. Genome-wide mapping and analysis of chromosome architecture. Nat. Rev. Mol. Cell Biol. 2016; 17:743–755

115. Li S, Tighe SW, Nicolet CM, et al. Multi-platform assessment of transcriptome profiling using RNA-seq in the ABRF next-generation sequencing study. Nat. Biotechnol. 2000;

116. Park PJ. ChIP-seq: advantages and challenges of a maturing technology. Nat. Rev. Genet. 2009; 10:669–680

117. Gamazon E, Huang RS, Dolan E, et al. Integrative Genomics: Quantifying Significance of Phenotype-Genotype Relationships from Multiple Sources of High-Throughput Data. Front. Genet. 2013; 3:

118. Ochs MF, Farrar JE, Considine M, et al. Outlier analysis and top scoring pair for integrated data analysis and biomarker discovery. IEEEACM … 2014;

119. Tyekucheva S, Marchionni L, Karchin R, et al. Integrating diverse genomic data using gene sets. Genome Biol. 2011; 12:R105

120. Shen R, Mo Q, Schultz N, et al. Integrative Subtype Discovery in Glioblastoma Using iCluster. PLOS ONE 2012; 7:e35236

121. Gevaert O. MethylMix: an R package for identifying DNA methylation-driven genes. Bioinforma. Oxf. Engl. 2015; 31:1839–1841

122. Witten DM, Tibshirani R, Hastie T. A penalized matrix decomposition, with applications to sparse principal components and canonical correlation analysis. Biostat. Oxf. Engl. 2009; 10:515–534

123. Fertig EJ, Markovic A, Danilova LV, et al. Preferential Activation of the Hedgehog Pathway by Epigenetic Modulations in HPV Negative HNSCC Identified with Meta-Pathway Analysis. PLOS ONE 2013; 8:e78127

124. Chen BJ, Causton HC, Mancenido D. Harnessing gene expression to identify the genetic basis of drug resistance. Mol. Syst. … 2009;

125. Andrews J, Kennette W, Pilon J, et al. Multi-Platform Whole-Genome Microarray Analyses Refine the Epigenetic Signature of Breast Cancer Metastasis with Gene Expression and Copy Number. PLOS ONE 2010; 5:e8665

126. Poisson LM, Taylor JM, Ghosh D. Integrative set enrichment testing for multiple omics platforms. BMC Bioinformatics 2011; 12:459

127. Liu Y, Devescovi V, Chen S, et al. Multilevel omic data integration in cancer cell lines: advanced annotation and emergent properties. BMC Syst. Biol. 2013; 7:14

128. Schulz MH, Pandit KV, Lino Cardenas CL, et al. Reconstructing dynamic microRNA-regulated interaction networks. Proc. Natl. Acad. Sci. U. S. A. 2013; 110:15686–15691

129. Yao L, Shen H, Laird PW, et al. Inferring regulatory element landscapes and transcription factor networks from cancer methylomes. Genome Biol. 2015; 16:

130. Fertig, Afsari B, Geman D. Learning Dysregulated Pathways in Cancers from Differential Variability Analysis. Cancer Inform. 2014; 61

131. Zhang Y, Liu T, Meyer CA, et al. Model-based analysis of ChIP-Seq (MACS) Genome Biol. 2008; 9 (9): R137. 2008;

132. Hennessey PT, Ochs MF, Mydlarz WW, et al. Promoter Methylation in Head and Neck Squamous Cell Carcinoma Cell Lines Is Significantly Different than Methylation in Primary Tumors and Xenografts. PLoS ONE 2011; 6:e20584

